# Modeling host-associating microbes under selection

**DOI:** 10.1101/2021.02.24.432736

**Authors:** Florence Bansept, Nancy Obeng, Hinrich Schulenburg, Arne Traulsen

## Abstract

The concept of fitness is often reduced to a single component, such as the replication rate in a given habitat. For species with complex life cycles, this can be an unjustified oversimplification, as every step of the life cycle can contribute to reproductive success in a specific way. In particular, this applies to microbes that spend part of their life cycles associated to a host, *i.e.* in a microbiota. In this case, there is a selection pressure not only on the replication rates, but also on the phenotypic traits associated to migrating from the external environment to the host and vice-versa. Here, we investigate a simple model of a microbial population living, replicating, migrating and competing in and between two compartments: a host and its environment. We perform a sensitivity analysis on the global growth rate to determine the selection gradient experienced by the microbial population. We focus on the direction of selection at each point of the phenotypic space, defining an optimal way for the microbial population to increase its fitness. We show that microbes can adapt to the two-compartment life cycle through either changes in replication or migration rates, depending on the initial values of the traits, the initial distribution of the population across the compartments, the intensity of competition, and the time scales involved in the life cycle versus the time scale of adaptation (which determines the adequate probing time to measure fitness). Overall, our model provides a conceptual framework to study the selection on microbes experiencing a host-associated life cycle.

## 1 Introduction

Fitness is a central concept in evolutionary biology, of particular importance for the theory of natural selection. Fitness measures how well a phenotype performs in terms of reproductive success, *i.e.* in terms of its ability to survive and reproduce. Natural selection, acting through reproduction and inheritance of the phenotypic traits, then leads to an increase in the population of the genotypes producing high fitness phenotypes [1].

In any system, fitness emerges mechanistically from birth and death events [2]. However, when it comes to the study of particular experimental systems or models, the question of how to measure fitness is often delicate, and fitness is often defined from the outset, as a phenomenological parameter. For example, fitness may be quantified as a net replication rate measured over a limited period of time in fixed laboratory conditions, or as a proportion of habitats successfully colonized. But none of these fitness components alone provides a holistic view of what fitness encompasses in natural conditions. Indeed, in nature, individuals undergo complex life cycles to produce new offspring, which makes fitness a multivariable function of all the life-history traits characterizing that organism’s life cycle. In addition to offspring production, this may include, for example, the ability of that offspring to migrate or disperse to the appropriate environments, or the ability to find mates in the case of sexual reproduction.

Life cycle complexity has been repeatedly shown to be important for the characterization of fitness. Historically, age-structured models have been developed to study human demography [3]. In the context of species conservation, or, at the other end of the spectrum, pest management, the focus has been on finding the “Achilles heels” of species life cycles to design efficient strategies to act upon them, in order to shape and preserve biodiversity [3]. This idea has further been developed theoretically, within the conceptual framework of metapopulation dynamics [4, 5]. Finally, life cycle complexity is also a concept central to the study of the onset of multicellularity, to understand why and how group replication can be selected for [6, 7].

The question of how life cycle components contribute to fitness is of particular relevance for the study of microbial communities that associate with hosts – microbiotas. Intricate life cycles are common in nature, where microbes can for example use hosts as vectors between different habitats [8, 9]. Having a living host as a habitat adds complexity to the assessment of fitness, given that the presence of the microbes may impact the host fitness and vice-versa. It is in fact the whole life cycle of host-associating microbes that is intertwined with the one of their host. Research has often been biased towards the host perspective, and has focused on how microbes can contribute to host fitness by extending the host functional repertoire, *e.g.* performing digestive or immune tasks [10, 11, 12]. An exception is epidemiology and parasitology, that have specifically addressed the impact of the host fitness on the pathogen, in the form of trade-offs between transmission and within-host virulence [13, 14, 15, 16]. But what about commensal relationships, where bacteria do not have a negative impact on the host fitness? In this context, what are the factors that determine a microbial population’s fitness?

Here, we propose a framework to assess the selection gradient acting upon the life history traits of a microbial population with a life cycle including host association. The gradient of selection determines the direction in the phenotypic space that evolution is expected to follow to maximize fitness. Our general aim is to provide a tool to compare the relative importance of the different life-history traits of a microbial population, starting only from the equations that describe the population dynamics experienced throughout the life cycle. We explore a simple continuous-time two-compartment model that allows microbes to migrate between a host and its environment. We use the method of sensitivity analysis [3] to infer how strongly the population growth rate depends on the traits we are considering. In the baseline version of the model, we consider unconstrained growth. Subsequently, we extend our framework numerically to include population size constraints.We define the local direction of the selection gradient as the optimal strategy for a microbial population to adapt to its life cycle, starting from the local values of the traits. We show the existence of defined regions of different optimal strategies in the phenotypic space in which it is either more beneficial to optimize growth or transmission. The boundaries of these regions are driven by modeling assumptions such as competition, and the probing time chosen to measure fitness.

## 2 Model

We focus on a single microbial type and ask how the growth rate of its whole population is affected by its life history traits. We study the population in two compartments corresponding to communicating habitats: the host and its environment. Let us write *n*_*H*_ for the number of host-associated microbes and *n*_*E*_ for the number of environmental ones. We define the life history traits of the microbial population as the rates at which individuals of the compartmental populations reproduce and die, compete and migrate from one compartment to another (Figure 1A). The net replication rates in the environment and within the host are *r*_*E*_ and *r*_*H*_, respectively. They could encompass both offspring production and death, and thus could be negative. The migration rates from the host to the environment and from the environment to the host are *m*_*E*_ and *m*_*H*_, respectively. We start with exponentially growing populations. We later introduce a competition of intensity *k*_*ij*_ experienced by the microbes of compartment *i* due to the abundance of microbes in the compartment *j*. We assume that the number of microbes is large enough to be described by differential equations and assume that all rates introduced above are constant.

**Figure 1:**
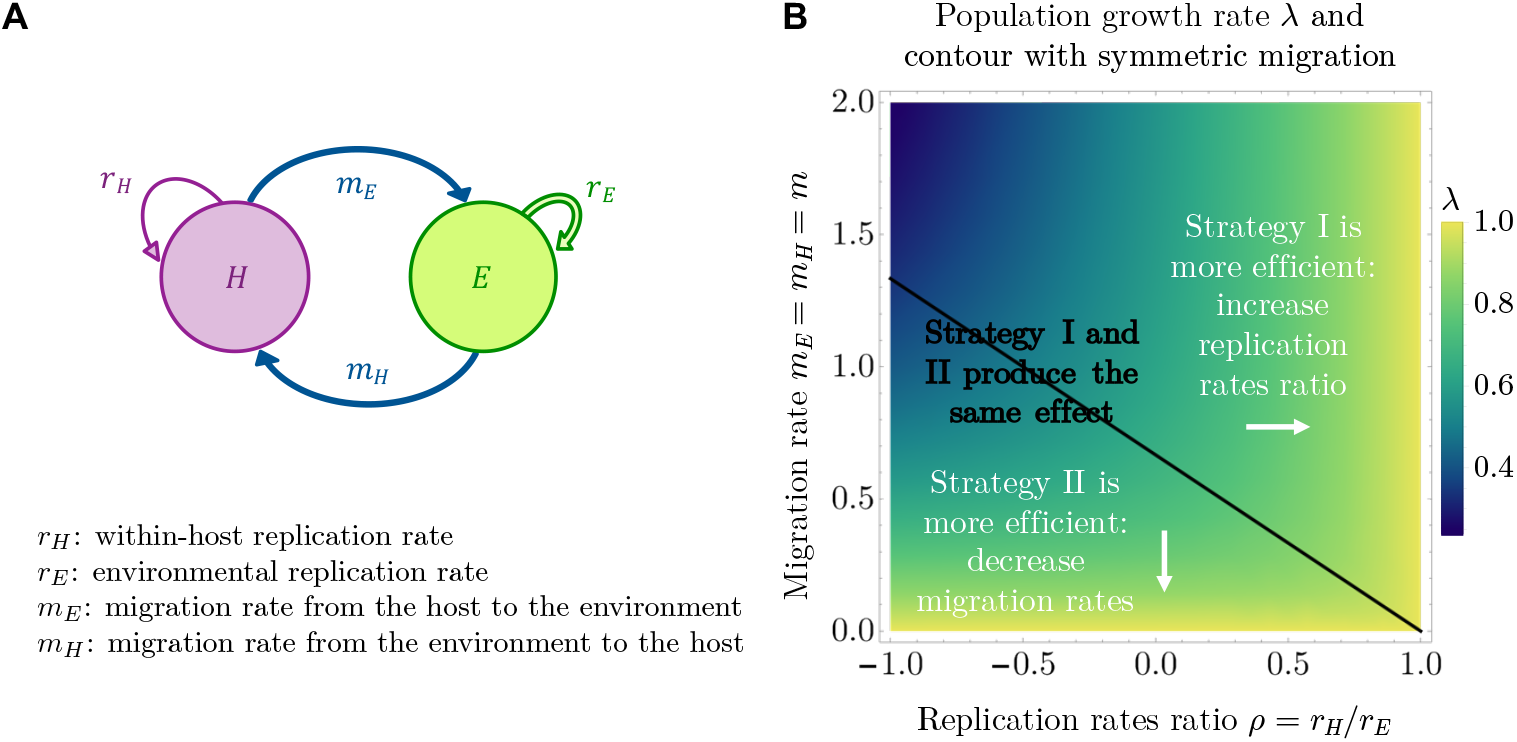
Optimal strategies in the baseline model (no competition). **(A)** Schematic diagram and definition of the rates for a microbial population migrating between a host and its environment and replicating in each compartment. For *r*_*E*_ > 0, the population increases exponentially and we ask how the exponential growth rate can be increased by changing the parameters of the model. **(B)** Population growth rate *λ* (color scale) on the trait space determined by *ρ* = *r*_*H*_/*r*_*E*_ and *m* = *m*_*H*_ = *m*_*E*_. The population growth rate *λ* is maximized for small *m* or for large *ρ*. In addition, we focus on sensitivities, which capture how strongly the population growth rate depends on the two traits. The contour line shows the line of the traits space that equalizes the absolute values of the two sensitivities derived analytically from equations 4 and 5, *|s*_*m*_*|*/*|s*_*ρ*_*|* = 1, delimiting the regions of optimality of the two different strategies. Note that we take the absolute values of the sensitivities, because in the baseline model the sensitivity of *λ* to increase in *m* is always negative, while it is always positive to increase in *ρ*. When *|s*_*m*_*|*/*|s*_*ρ*_*|* < 1, the optimal strategy is to increase the replication rates ratio (strategy I). When *|s*_*m*_*|*/*|s_ρ_|* > 1, the optimal strategy is to decrease the migration rate (strategy II).

This leads to the general equations

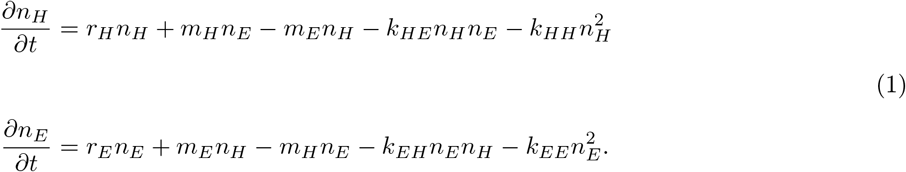

In the following, we first consider unconstrained growth, where there is no competition (*k*_*EE*_ = *k*_*HH*_ = *k*_*EH*_ = *k*_*HE*_ = 0), before adding global competition (*k*_*EE*_ = *k*_*HH*_ = *k*_*EH*_ = *k*_*HE*_ = k), competition limited to one of the compartments (*k*_*EH*_ = *k*_*HE*_ = 0 and *k*_*EE*_ ≠ 0 or *k*_*HH*_ ≠ 0), and finally, equal competition in each of the compartments (*k*_*EH*_ = *k*_*HE*_ = 0 and *k*_*EE*_ = *k*_*HH*_ = k). While in nature it is likely that none of the *k*_*ij*_ vanishes and that a wide range of values are possible, the study of these limit cases gives powerful insights into what is to be expected in a wide range of situations.

## 3 Results

### 3.1 Baseline model: no competition

We start by assuming no competition and consider unconstrained growth in each of the two compartments. In this case, the equations describing our model become linear and can be rewritten in matrix form [3] as

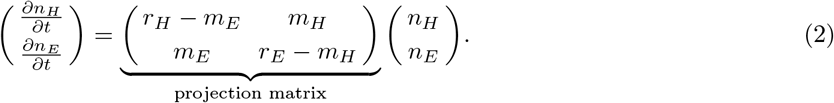

The dominant eigenvalue *λ* of the above-defined projection matrix gives the asymptotic growth rate of the whole population. This quantity is an appropriate measure of fitness [3] insofar as it measures reproductive success and recapitulates the effects of all the life history traits. The dominant right eigenvector represents the stable distribution in the two compartments of an exponentially growing population. The value of *λ* can be calculated at each point of the phenotypic space defined by the ranges of possible values that could be taken by the life-history traits *r*_*E*_, *r*_*H*_, *m*_*E*_, and *m*_*H*_. The dependence of *λ* on these traits tells us at which points of the phenotypic space fitness is maximized and how it can be increased at all other points.

From the projection matrix, we calculate the dominant eigenvalue as

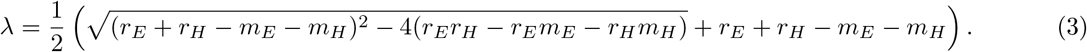

Note that if microbes replicate at the same rate in the host and in the environment, *i.e.* if *r*_*E*_ = *r*_*H*_ = *r*, *λ* simplifies to *r*, regardless of the migration rates *m*_*H*_ and *m*_*E*_. When there is an asymmetry between the two replication rates however, which is very likely to be the case in nature, then the migration rates also affect the population growth rate. In the following sections, we study this effect compared to the effect of the replication rates. We arbitrarily set *r*_*H*_ ≤ *r*_*E*_, and *r*_*E*_ > 0 – otherwise the population gets extinct. In biological terms, this corresponds to the situation where the microbial population is initially more adapted to the environment than to the host and thus grows faster in the environment. But mathematically, in this model, host and environment are symmetrical, *i.e.* they only differ by the rates defined above. Thus, the chosen direction of this inequality does not carry any strong meaning, and there is no loss of generality in making this choice. In particular, one can access the opposite biological situation where microbes replicate faster in the host than in the environment – as is the case for viruses, that can only replicate in the host (*r*_*H*_ > 0) but decay in the environment (*r*_*E*_ < 0) – by a single switch of the index *E* and *H*.

Let us first study the case where the migration rates from and towards the environment are equal, *i.e. m*_*E*_ = *m*_*H*_ = *m* > 0. Let us denote 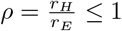 the ratio of the replication rates. Then, setting *r*_*E*_ =1 to scale time (and thus, measuring all other rates in units of the replication rate of the microbe in the environment), *λ* reduces to

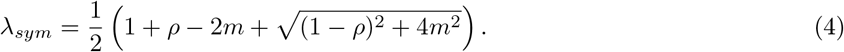

For any fixed positive value of *m*, *λ*_*sym*_ is a strictly increasing function of *ρ*, which reflects the fact that increasing *ρ* allows for additional growth within the host. We will limit ourselves to the study of *ρ* ≥ −1, which guaranties a positive value for *λ*_*sym*_. For any fixed value of *ρ*, *λ*_*sym*_ is a decreasing function of m, which reflects the fact that for increasing *m*, microbes are increasingly lost towards the host, where growth is slower than in the environment. Figure 1B shows the value of *λ*_*sym*_ on the reduced phenotypic space defined by *ρ* and m. The maximum possible value for *λ* is 1 (in units of *r*_*E*_). This value is achieved either by increasing the ratio of replication rates between host and environment, so that both microbial populations grow at the same rate (strategy I), or by reducing migration between host and environment (strategy II). This second strategy allows microbes to spend a longer time in the environment on average. Note however, that this strategy is limited, since setting *m* to zero decouples the two compartments completely, in which case the two subpopulations grow independently at different rates.

How strong is the selection on these traits? This question can be approached by inferring how strongly the population growth rate depends on the traits we are considering. One standard approach to measure this is sensitivity analysis [3]. One defines the sensitivity of the population growth rate *λ* achieved by the phenotype described by the vector **x**= (*x*_1_,…, *x*_*N*_) in the trait space to its i^*th*^ life-history trait as

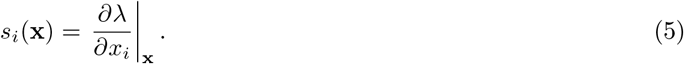

This quantity gives the change in the value of *λ* that results from a small increment of the trait *i*. It is a local property that can be calculated for each point **x** of the trait space. The vector of the sensitivities at point **x** gives the direction of the selection gradient on the fitness landscape. In other words, to achieve effcient phenotypic adaptation, the population should move in the trait space following the direction of this gradient.

If the population can invest in phenotypic adaptation only by tuning one of its life-history traits at a time, then it should act upon the trait that has the largest (absolute) sensitivity at the current position of the population in the trait space. This reasoning allows to divide the trait space into regions of distinct optimal strategies, as shown in Figure 1B. In the regime of high migration rates (*i.e.* when the switch between the compartments is so rapid that the population is almost experiencing a habitat having average properties between the host and the environment), strategy I (increasing *ρ*) becomes almost always optimal, except for small replication ratios, where there is almost no replication in the host. In summary, migration rates are important when replication in the host is slow compared to the environment, and when migration itself is slow. These conclusions remain qualitatively unchanged with asymmetric migration rates, as discussed in more detail in the electronic supplementary material (ESM) section A.1.

### 3.2 Model with global competition between all microbes

In the baseline model, there are no constraints on population growth. In nature, however, microbial populations do face limits to their growth. Since the equations above are linear and can only give rise to exponential growth or exponential decay, they can only describe the dynamics of a population over a limited period of time. In order to account for saturation and competition during growth, we thus need to introduce non-linear terms to the equations 1. The study of this kind of systems often focus on long term dynamics, yet it can be of high practical relevance to study the transient optimal strategies, as shorter timescales are often relevant in the real world – whether it be due to experimental constraints or to ecological disturbances and perturbations [17]. Since we are going to consider some out-of equilibrium dynamics, in particular in the section with competition limited to one of the compartments, and because we are also interested in transient properties, we will adopt a numerical approach based on population sizes [18, 19].

In this section we study the case of a microbial population limited in size by global competition occurring at rate *k* = *k*_*HH*_ = *k*_*EE*_ = *k*_*EH*_ = *k*_*HE*_. This situation could correspond to a microbiota living in direct contact with an external environment, *e.g.* on the surface of an organism. Alternatively, what we call the “environment” in our model could represent another host compartment in direct contact with the other, like the gut lumen and the colonic crypts. In that case, microbes living in association with the host are in direct contact with those in the environment and can mutually impact each other’s growth. This is of particular relevance if both microbial subpopulations rely on and are limited by the same nutrients for growth.

From the microbial abundances in the different compartments obtained by numerically solving the equations, one can build a proxy for the population growth rate. To remain coherent with the previous section, we define

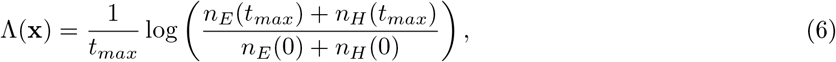

*i.e.* the effective exponential growth rate of the whole microbial population – which captures the individual level reproductive success, since we consider a homogeneous population – over the chosen period of time [0, *t*_*max*_]. There are several fundamental differences between the effective exponential growth rate Λ in a non-linear system and the population growth rate *λ*; in a linear system, the dominant eigenvalue of the projection matrix as defined in the baseline model. First, Λ only provides a measure of growth for the whole population, but does not correspond to the asymptotic growth rate of each subpopulations as it was the case with *λ* in the baseline model. In fact, it is not either the asymptotic growth rate of the whole population: in the case of global saturation, replication stops when the carrying capacity is reached, and the asymptotic growth rate for the whole population is thus zero. Therefore, the choice of the probing time *t*_*max*_ has an impact on Λ, which we will see in more detail below. Second, the choice of the exact form of Λ now implies biological assumptions on the selection pressure felt by the population: choosing the effective exponential growth rate over the whole population as we do implies that selection is acting on the whole population evenly. There may be some situations, for example experiments in which the population of one of the compartments is artificially selected for, where it would make more sense to define Λ as the effective exponential growth rate over just this subpopulation. This may lead to different conclusions, in particular at the transient scale. One must thus adapt Λ to the specifics of the modeled system. In addition, the choice of *t*_*max*_ itself has a biological meaning, and should in particular not exceed the time upon which the dynamics of the system are accurately described by the set of equations. This may also be determined by experimental times.

We now calculate the sensitivity of Λ in the direction of the trait *i* at the point **x** of the phenotypic space as

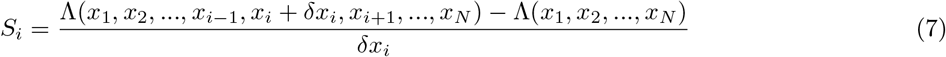

with *δx*_*i*_ the discretization interval, and *N* the number of traits defining a phenotype **x**.

For the numerical approach, additional choices need to be made. First, the trait space needs to be discretized. Then, to calculate Eq. 7, one needs to choose a set of initial conditions and a probing time at which to measure the population sizes, as exposed in details for the linear case in [17]. Finally, we need to choose the discretization interval *δx*_*i*_. In the following, we always choose *δx*_*i*_ sufficiently small for convergence, *i.e.* so that it does not significantly impact the numerical values of the sensitivities, and focus on the choices of the other parameters (probing time and initial conditions) and the influence of the competition intensity *k*. One strategy to explore the possible impact of initial conditions is to use “stage biased vectors” [17], *i.e.* extreme distributions of the population. This corresponds to initial conditions where microbes either exist only in the host or only in the environment.

In Figure 2, we show how the contour lines delimiting the two optimal strategies change with the final time *t*_*max*_ chosen to measure the population growth rate and with the intensity of competition *k*, for these two extreme cases: *n*_*E*_(0) = 0, *n*_*H*_ (0) = 1 and *n*_*E*_(0) = 1, *n*_*H*_ (0) = 0. In all cases, with sufficiently long *t*_*max*_, the contours converge to the contour plot of the baseline model in the previous section. This is expected, since competition here affects all the microbes in the same way, so that the equilibrium distribution is the same as the asymptotic distribution of the baseline model (given by the dominant eigenvector). Mathematically, global competition can be seen as a modification of the baseline projection matrix by subtracting an identity matrix times a scalar depending on time. This does neither affect the eigenvectors nor the dependence of the dominant eigenvalue on the traits.

**Figure 2:**
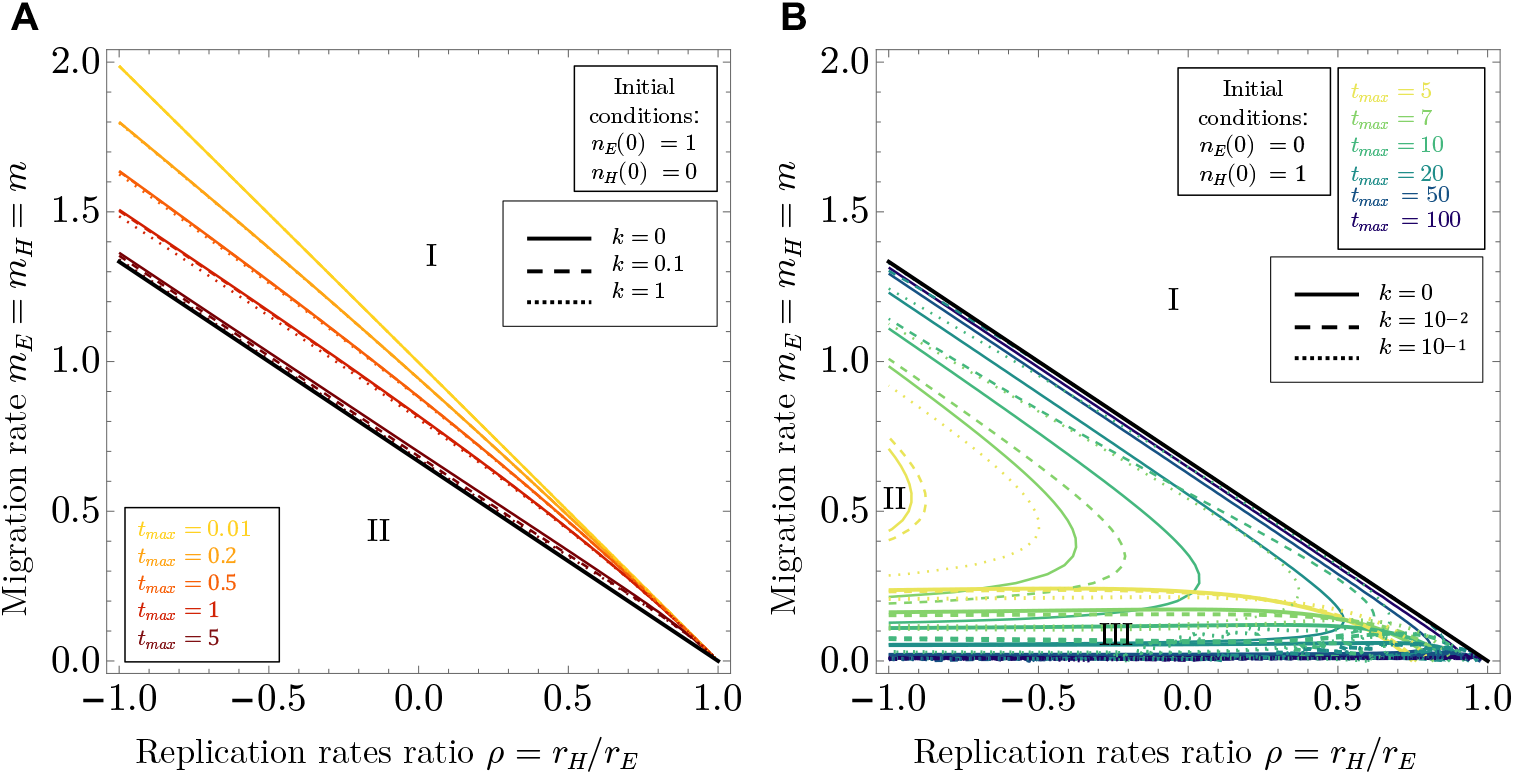
Optimal strategies in the model with global competition. **(A)** Change in the contour line delimiting the regions of optimality of the two strategies (strategy I: increasing the replication rates ratio; strategy II: decreasing migration) with *t*_*max*_, the time chosen to measure the final population size, measured in units of 1/*r*_*E*_. Initially all the microbes are in the environment. Because in this model all the microbes are equally impacted by competition, with *t*_*max*_ large enough, one recovers the contour line of the baseline model calculated analytically (black line). Continuous lines: *k* = 0, *i.e.* no competition. Dashed lines: increasing values of *k* (competition intensity). **(B)** Extension of A with all the microbes in the host initially. In this case the convergence to the case without competition appears to be slower, while increasing the value of *k* seems to accelerate it. A third optimal strategy (III: increasing migration) appears around *m* = 0, delimited from strategy I by thicker lines.

In the case where all the microbes are initially in the host (Figure 2B), the convergence to the baseline case requires higher values of *t*_*max*_ than in the case where all the microbes are initially in the environment (Figure 2A). Intuitively, this corresponds to the fact that convergence to the baseline distribution requires population growth, and growth is slower if all the microbes are initially in the host compartment – where replication is slower. There is thus a time delay between these two situations, corresponding to the time it takes for migration to carry microbes into the environment – where replication is faster. When all the microbes are initially in the host (Figure 2B), we also observe the appearance of a third optimal strategy around *m* = 0: increasing the migration rate. In this unfavorable condition (*m* = 0 and an initially empty environment), increasing the microbial flux towards the environment becomes more important than limiting the flux of microbes leaving it (which is nonexistent when *m* = 0). At large *t*_*max*_ there is a direct transition between strategy II and strategy III when increasing m from zero, thus the two contour lines overlap.

Finally, we observe that the intensity of competition has only a small effect on the contours when all the microbes are initially in the environment but a larger effect when all the microbes are initially in the host. In both cases, adding competition (*k* > 0) appears to accelerate convergence to the baseline contour. This is because during the transient dynamics, the distribution balances to the expected asymptotic distribution from the initial conditions. In most cases, this equilibrium distribution is a mixed state, where a part of the population lives in the environment and another in the host. To reach this balance quickly from a pure biased state necessitates immigration to and replication in the initially empty compartment, while immigration to and replication in the other compartment remain slow. Competition limits the growth in the compartment that is not initially empty, and thus helps this balancing process. This effect is even stronger if the initially empty compartment happens to be the environment, where microbes replicate faster. This explains why the effect of k is stronger in this case, see Figure 2B.

### 3.3 Model with competition within one of the compartments only

In this section we consider competition happening inside one of the compartments only (*i.e. k*_*EH*_ = *k*_*HE*_ = 0 and *k*_*EE*_ ≠ 0 or *k*_*HH*_ ≠ 0). We will start by considering competition in the host only, as it seems likely, from the biological point of view, that resources may be more limited in the host than in the environment. However, in a second step we also look at the case with competition limited to the environment. Even if this situation may seem less likely at first sight, one should bear in mind that it also covers the case of competition limited to a host where replication is faster than in the environment (*r*_*H*_ > *r*_*E*_), provided a switch of the *H* and *E* index.

In the case where competition is limited to only one of the compartments, we do not expect an equilibrium population size to be reached for all traits combination of the phenotypic space. If migration is not sufficiently important, the subpopulation in the unconstrained compartment keeps growing exponentially faster than the other subpopulation, which contribution to the global population thus becomes rapidly negligible. At sufficiently high migration rates however, an equilibrium is expected, because the microbes switch habitats sufficiently rapidly for competition to be globally effective, although it directly affects only one of the compartments.

#### 3.3.1 Competition in the host only (slow-replicating compartment)

When there is competition in the host only, there is no (positive) equilibrium for all *m* < 1. In this case, replication inside the host should have less importance because the host subpopulation size becomes negligible compared to the one of the environment. On this region of the phenotypic space we thus expect the sensitivity of parameter *ρ* to tend to zero with increasing probing times *t*_*max*_, and the contour lines to be shifted to increase the area of optimality of strategy II, whatever be the other parameters (initial conditions, intensity of competition).

Figure 3 verifies this verbal argument: as expected, for a fixed *t*_*max*_, we recover the shape of the fitness landscape of the baseline model for small values of *k*. Strategy I (increasing the replication rates ratio) however, sees its area of optimality reduced with increasing values of *k* (Figure 3A). The values of Λ also become smaller overall: growth is slower due to competition.

**Figure 3:**
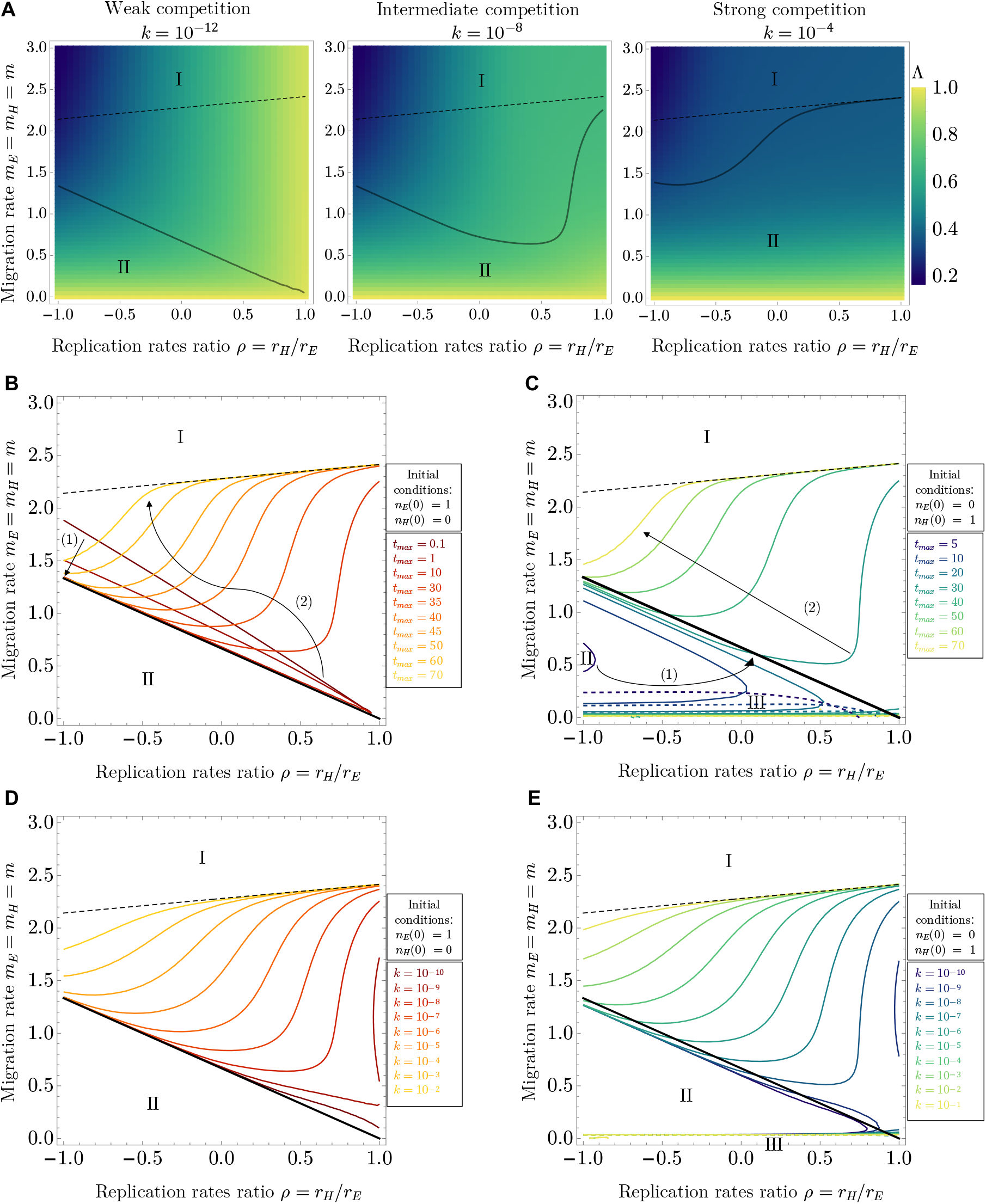
Optimal strategies in the model with competition in the host only. **(A)** Change in the fitness landscape with the within-host competition intensity *k* = *k*_*HH*_. Black line: contour of equal sensitivities. Thin dashed line: contour of equal sensitivities of the equilibrium population sizes. Other parameters: *t*_*max*_ = 30, *n*_*E*_(0) = 1, *n*_*H*_ (0) = 0. **(B-E)** Change in the contour lines delimiting the regions of optimality of the strategies with **(B-C)** *t*_*max*_ (probing time chosen to measure the population size) and **(D-E)** *k* (within-host competition intensity), with initial conditions where all the microbes are in the environment (B and D, *n*_*E*_(0) = 1, *n*_*H*_(0) = 0) or where all the microbes are initially in the host (C and E, *n*_*E*_(0) = 1, *n*_*H*_ (0) = 1). Solid lines: limit between the regions of optimality of strategy I (increasing the replication rates ratio) and II (decreasing migration). Dashed lines: between strategies I and III (increasing migration). Other parameters: *k* = 10^−8^ (B-C), *t*_*max*_ = 30 (D-E). In the case of competition limited to the host, whatever be the initial conditions, at sufficiently large *t*_*max*_, or sufficiently large *k*, the region of optimality of strategy I (increasing the replication rates ratio) tends to narrow down and shift out of the *m* < 1 region to reach the contour of equal sensitivities of the equilibrium population sizes (thin dashed line). The arrows in panels (B) and (C) indicate the two steps of contour shift with increasing *t*_*max*_: in the first phase, when competition has a limited effect due to low abundances, the contours approach the limit of no competition (black line, as in Figure 2). In the second phase, competition in the host kicks in, and the contours move away from the baseline limit.

As expected, for a fixed value of *k* = *k*_*HH*_, the contours delimiting the two optimal strategies shift to reduce the area of optimality of strategy I with large values of *t*_*max*_ (Figure 3B-C). The disappearance of the contours from the region where *m* < 1 takes place in two steps. First, with small *t*_*max*_ values, the effect of competition is not yet apparent due to the low initial abundance of microbes, and the contours start by getting closer to the reference contour of the baseline model, just as was observed in Figure 2. In the second step, with larger *t*_*max*_, the effect of competition becomes apparent and the contours are shifted out of the *m* < 1 region, ultimately reaching a close-to horizontal limit which can be calculated analytically by performing sensitivity analysis on the equilibrium population sizes.

Additionally, when initially the microbes are in the host (Figure 3C), for the same reasons as in the previous section, we can again observe the appearance of the third strategy, increasing the migration rate, around m = 0.

The impact of increasing competition *k* at fixed *t*_*max*_ on the contour lines delimiting the two optimal strategies is clear in Figure 3D and E. We see that increasing *k* and increasing *t*_*max*_ have very similar effects: with small values of *k*, the baseline case is recovered, as expected. When increasing *k* sufficiently, the contour line is shifted out of the *m* < 1 region with strategy I, *i.e.*increasing the replication rates ratio, until reaching the equilibrium limit. This is because increasing growth in the host can only have a limited effect when growth in the host is limited by competition, which makes strategy II comparatively more effcient.

#### 3.3.2 Competition in the environment only (fast-replicating compartment)

When there is competition in the environment only, there is no (positive) equilibrium for all *m* < *ρ*. In this region of the phenotypic space, the size of the environmental population becomes substantially smaller than that of the host-associated population after some time. As a consequence, strategy I (increasing the replication rate within the host) becomes more important, so that we see its area of optimality extend, see Figure A.2. For a fixed *t*_*max*_, with a small value of *k* we recover the shape of the fitness landscape from the baseline model with no competition, but increasing *k* shifts the contour line to lower *ρ* until the strategy II (decreasing migration) disappears from the *m* < *ρ* region and the delimitation of the strategies approaches the contour of equal sensitivities of the equilibrium population sizes, calculated analytically. Like in the previous sections, we also observe the appearance of a third optimal strategy around *m* = 0, increasing migration. Unlike in the previous sections, this time the third strategy also appears when the microbes are all initially in the environment (Figure A.2B and D), and is also predicted by the sensitivity analysis of the equilibrium population sizes. Intuitively, having competition in the fast-replicating environment reduces the advantage of starting with a microbial population exclusively located there, and in this case too migration towards the host becomes initially more important than limiting migration out of the environment. For a fixed value of *k*, with increasing *t*_*max*_ the contour line starts by getting closer to the baseline model contour, before diverging from this limit and approaching the contour of equal sensitivities of the equilibrium population sizes. This finally leaves strategy I as the only optimal strategy on almost all the phenotypic space at sufficiently long times (Figure A.2B-C), except for a region of small *ρ* and intermediate *m*. Increasing *k* for a fixed value of *t*_*max*_ (Figure A.2D-E) has a very similar effect on the contour, except for the initial dynamics towards the baseline model.

### 3.4 Competition of same intensity in each compartment

When there is competition of equal intensity in the host and the environment (*i.e. k*_*EH*_ = *k*_*HE*_ = 0 and *k*_*EE*_ = *k*_*HH*_ = *k*), we observe very similar results to the previous section, with competition in the environment only (see Figure 4): increasing *k* or increasing *t*_*max*_ leads to the disappearance, at long times, of the area of optimality of strategy II (decreasing migration), except for a distinct region of small *ρ* and intermediate *m*, predicted by the contour of equal sensitivities of the equilibrium population sizes. This implies that the effect of competition in the fast-replicating compartment has a dominating effect on the global population growth rate.

**Figure 4:**
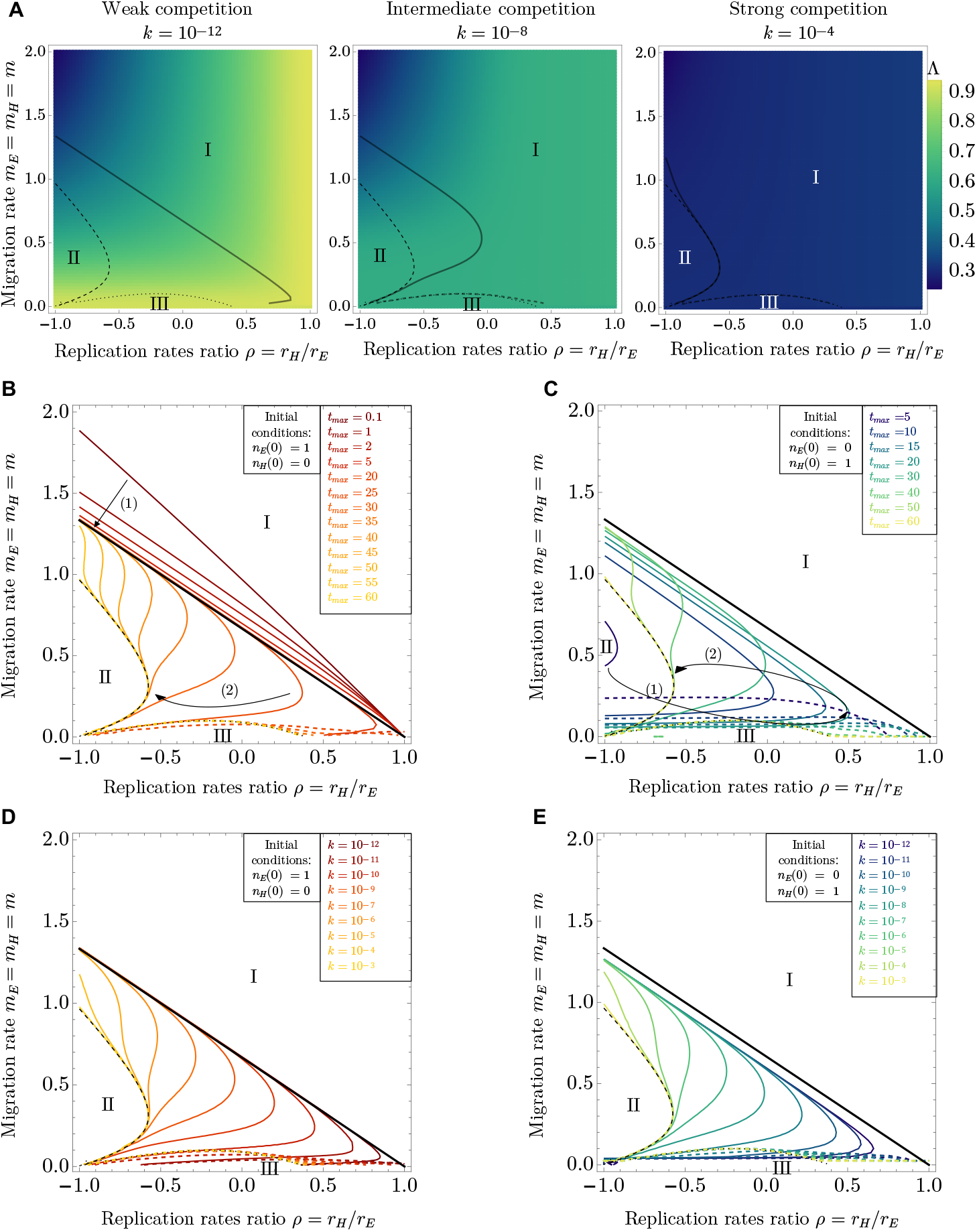
Optimal strategies in the model with limited growth in the host and the environment. **(A)** Change in the fitness landscape with the competition intensity *k*_*HH*_ = *k*_*EE*_ = *k*. Black thick lines: contours of equal sensitivities (solid line: between strategy I and II, dashed line: between I and III). Black thin lines: contours of equal sensitivities of the equilibrium population sizes (dashed: between I and II and dotted: between I and III). Other parameters: *t*_*max*_ = 30, *n*_*E*_(0) = 1, *n*_*H*_ (0) = 0. **(B-E)** Change in the contour lines delimiting the regions of optimality of the strategies with **(B-C)** *t*_*max*_ (time chosen to measure the population size) and **(D-E)** *k* (competition intensity), in the case of initial conditions where all the microbes are in the environment (B and D, *n*_*E*_(0) = 1, *n*_*H*_ (0) = 0) or in the case where all the microbes are initially in the host (C and E, *n*_*E*_(0) = 0, *n*_*H*_ (0) = 1). Solid lines: limit between the regions of optimality of strategies I (increasing the replication rates ratio) and II (decreasing migration), and dashed lines: between I and III (increasing migration). Other parameters: *k* = 10^−8^ (B-C) and *t*_*max*_ = 30 (D-E). In the case of competition in the host and in the environment, the effect of the competition in the environment (the fast-replicating compartment) seems to dominate, so that at sufficiently large times, the region of optimality of strategy II (decreasing migration) is reduced, and its limit approaches the contour of equal sensitivities of the equilibrium population sizes (black dashed line: between I and II). The black dotted line shows the contour of equal sensitivities of the equilibrium population sizes delimiting strategies I and III. The arrows in panels (B) and (C) indicate the two steps of contour disappearance with increasing *t*_*max*_: in the first phase, when competition has a limited effect due to low abundances, the contours approach the limit of no competition (black solid line, as in Figure 2). In the second phase, competition kicks in and the contours move away from the baseline limit.

## 4 Discussion

Out in the wild, microbes experience complex life cycles. Each of their steps can contribute to the overall reproductive success. In general, microbial fitness is thus more complex than the common approximation of growth yield used in the lab. This is particularly true for microbes experiencing life cycles that involve only a limited phase of host association, which translates as a selection pressure on phenotypic traits associated to migrating from an external environment to the host and vice-versa. A framework to study fitness in all its complexity is needed in the field of microbiome studies, which could benefit from approaches first introduced in demography. Here, we investigate a model of a microbial population living, replicating, migrating, and competing in and between two compartments: a host – assumed to be, throughout the paper, a compartment where replication is slower – and its environment. To analyze the selection gradient experienced by the microbial population going through this biphasic life cycle – with phases in the environment and phases in the host – we perform sensitivity analysis. We focus on the leading direction of the selection gradient at each point of the phenotypic space, thereby defining an optimal strategy for the microbial population to maximize its fitness.

We show that in the case of unconstrained exponential growth in both the compartments, there are two optimal strategies: increasing the replication rate in the host compared to the environment (strategy I), and decreasing the migration rates (strategy II) to maximize the time spent in the fast-replicating compartment. The first strategy is optimal at initially high ratios of replication rates and high migration rates, while the second strategy is optimal at initially small migration rates and small ratio of replication rates.

Next, we extend the model to a scenario where microbial growth is limited by competition. We start with global competition, a case which could describe competition for a resource homogeneously shared between the host and the environment. Biologically, this corresponds to communities of microbes that are associated with hosts, i.e. microbiotas, but have extensive contact with the environment, as the skin or other epithelial microbiotas for example [20, 21]. In this case, we show that apart from a transient effect, the optimality of the strategies is conserved from the case without competition. With competition in the host only (the slow-replicating compartment), at longer probing times, or at higher competition intensities, the strategy I (increasing the ratio of replication rates) is disfavored when migration out of the environment is slower than replication in the environment, *i.e.* where there is no equilibrium. Strategy II (decreasing the migration rates) thus increases its area of optimality. Inversely, with competition in the environment only (the fast-replicating compartment), or with competition of same intensity within the host and within the environment, the strategy II is selected against when migration out of the host is slower than replication in the host, leaving strategy I as the only optimal strategy on this region of the parameter space. Unsurprisingly, this suggests that competition within the fast-replicating compartment dominates the effect on the selection gradient.

While this analysis provides crucial information on the selection gradient that shapes microbial adaptation to life cycles involving host association, it does not take into account the evolvability of the traits themselves. Although the selection gradient is a good indicator of the expected evolutionary path in the phenotypic space, the underlying genotype/phenotype mapping does not always allow for this path to be taken [22, 23, 24, 25], and the outcome of evolution may thus be different. The discrete nature, the non-additivity and non-linearity of genetic information, as well as the existence of costs, trade-offs and evolutionary constraints may prevent the predicted continuous change on the phenotypic trait. In addition, using sensitivities is built on the assumption that adaptation generates additive changes in life history traits. Although this is a common assumption, different choices are sometimes made. For example, multiplicative changes of the traits are assumed in elasticity analysis [3, 18, 24, 26], which presents the advantage of manipulating only proportional changes and thus non-dimensional quantities, but deals poorly with traits that can take the value of zero. These fundamental assumptions can sometimes result in different inferred selection gradients, as was shown for example in the context of age-classified populations [27]. However, although the exact shapes of the contours are modified, we have checked that our qualitative results remain robust when applying elasticity instead of sensitivity analysis.

Stepping back, we can evaluate the predictions of our model in the light of biological observations. Evolution experiments where microbial populations are serially passaged through a host and an environment are of particular interest here, to assess the response to selection resulting from biphasic life cycles. The key role of microbial immigration during the initial adaptation to their zebrafish host has for example been highlighted in [28]. In *Drosophila* [29] and in *C. elegans* [30], experimental selection towards host association resulted in adaptive changes in microbial life history with a direct impact on host fitness. In detail, in the first case, there is evolution towards by-product mutualism, and in the second, which concerns an initially pathogenic population, evolution towards less virulence and an increased carrying capacity.

Conceptually, using the effective population growth rate as a measure of fitness provides a complementary insight to invasion fitness approaches [31, 32] developed to analyze such evolution experiments, for example in [33, 34]. While invasion fitness analysis relies on assessing the long term chances of successful invasion of an established population at equilibrium by a new mutant strain of defined traits values, sensitivity analysis of the effective population growth rate provides a systematic framework that can be applied to out-of-equilibrium systems, and provides information on shorter time scales. Both frameworks rely on different proxies to assess a fitness capturing its different components - in one case, the frequency of patches where the microbe is present, and in the other, the microbial population growth, but both frameworks converge on the key role of migration between compartments. In fact, in many common cases like global competition, the long-term predictions of invasion fitness are recovered with the sensitivity analysis of the effective growth rate by setting *t*_*max*_ sufficiently large [18].

In future work, our framework could be extended in different directions to capture additional characteristics of microbial life cycles in host association. The first extension could be to increase the number of compartments. While the question of fluctuating environments has been studied before, in discrete times or in a different context [7, 18], in our context it may be profitable to consider and include host population dynamics. This would notably allow us to include microbial traits that affect host fitness in our analysis. A second direction could be to include stochasticity and non-homogeneities. Indeed, our deterministic description is valid only if the size of the microbial population is sufficiently large at all times and can only describe the average selection gradient experienced by the population. Introducing stochasticity would allow the study of differentiation, which may play a role in the response to complex life cycles. Differentiation, in the form of speciation, phenotypic plasticity, or bet-hedging is indeed observed in evolution experiments and natural microbial populations [35, 36, 37, 38, 39, 40]. It is also observed in host-associated populations [41] and may thus be expected in evolution experiments that include a host-association phase. Finally, a key aspect that we have so far excluded is spatiality. Effects of spatiality on the selection gradient are known for example in a simple Petri dish system, where the existence of an optimal expansion speed for a given habitat size is shown [42, 43]. Generally, hosts are highly structured habitats with variation in nutrients and chemical and physical gradients shaping for example the gut [44, 45, 46], which may also favor differentiation. The introduction of several compartments or sub-compartments within the hosts could represent a first step in this direction.

Thus, our model provides the key ingredients to study the consequences of host association for a microbe. It meets the need to conceptualize fitness as a holistic measure that captures all the aspects of microbial life cycles. With the development of this framework, we aim to contribute to a better understanding of the mutual benefits that microbes and hosts can retrieve from such associations.

## Acknowledgments

The authors thank the Evolutionary Theory Department in the MPI Ploen for useful feedback and discussions, and Stefano Giaimo, Roman Zapién-Campos and Claude Loverdo for careful reading of an earlier version of the manuscript. All authors acknowledge funding and support from the CRC 1182: Origins and Functions of Metaorganisms, project A4.

