## Supplementary material for "Modeling host-associating microbes under selection"

### A Supplementary material

#### A.1 Asymmetric migration rates

Let us now consider an asymmetry between the migration rates. Setting $m_E = m$ and $m_H = \alpha m_E$, the expression of the population growth rate becomes

$$\lambda_a = \frac{1}{2}\left(1 + \rho - (1+\alpha)m + \sqrt{(m(1+\alpha) - 1 - \rho)^2 - 4(\rho - m(1+\alpha\rho))}\right). \tag{8}$$

The resulting shape of the fitness landscape is qualitatively unchanged: $\lambda$ is still maximized for low values of the migration rates and high values of the replication rates ratio, and the maximum value for the population growth rate $\lambda$ is still 1 (in units of $r_E$). However, the depth of the landscape changes with $\alpha$: the larger $\alpha$ (*i.e.* the more migration towards the host is favored compared to migration into the environment), the larger the amplitude of the landscape, and the steeper the slope to climb to reach the maxima. How does this translate in terms of optimal strategy? Figure A.1 shows how the contour delimiting the optimality of the two strategies is modified. When $\alpha$ is increased the range of optimality of strategy II is narrowed. This is expected, as the most important factor to maximize $\lambda$ is to stay in the environment – where replication is faster – as long as possible, and thus to reduce $m_H$. In the unfavorable condition of a large $\alpha$, microbes need to reduce $m$ more to achieve the same effect on $m_H$. Thus, strategy II (decreasing the migration rates) becomes less efficient.

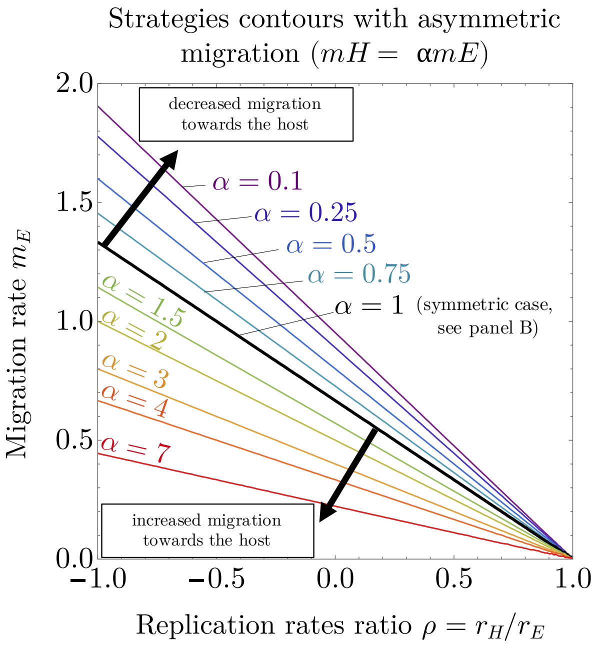


Figure A.1: Change in the contour line delimiting the regions of optimality of the two strategies defined in panel B with $\alpha$, the ratio of the migration rates. The black thick line shows $\alpha = 1$, *i.e.* the symmetric case identical to the contour shown in panel B. Increasing the migration towards the host is unfavorable for the considered population, because of the assumption that $r_H \leq r_E$. With high values of $\alpha$, one needs to decrease $m$ more to achieve similar results on $\lambda$, thus decreasing the area of optimality of strategy II.

#### A.2 Competition in the environment only: supplementary figure

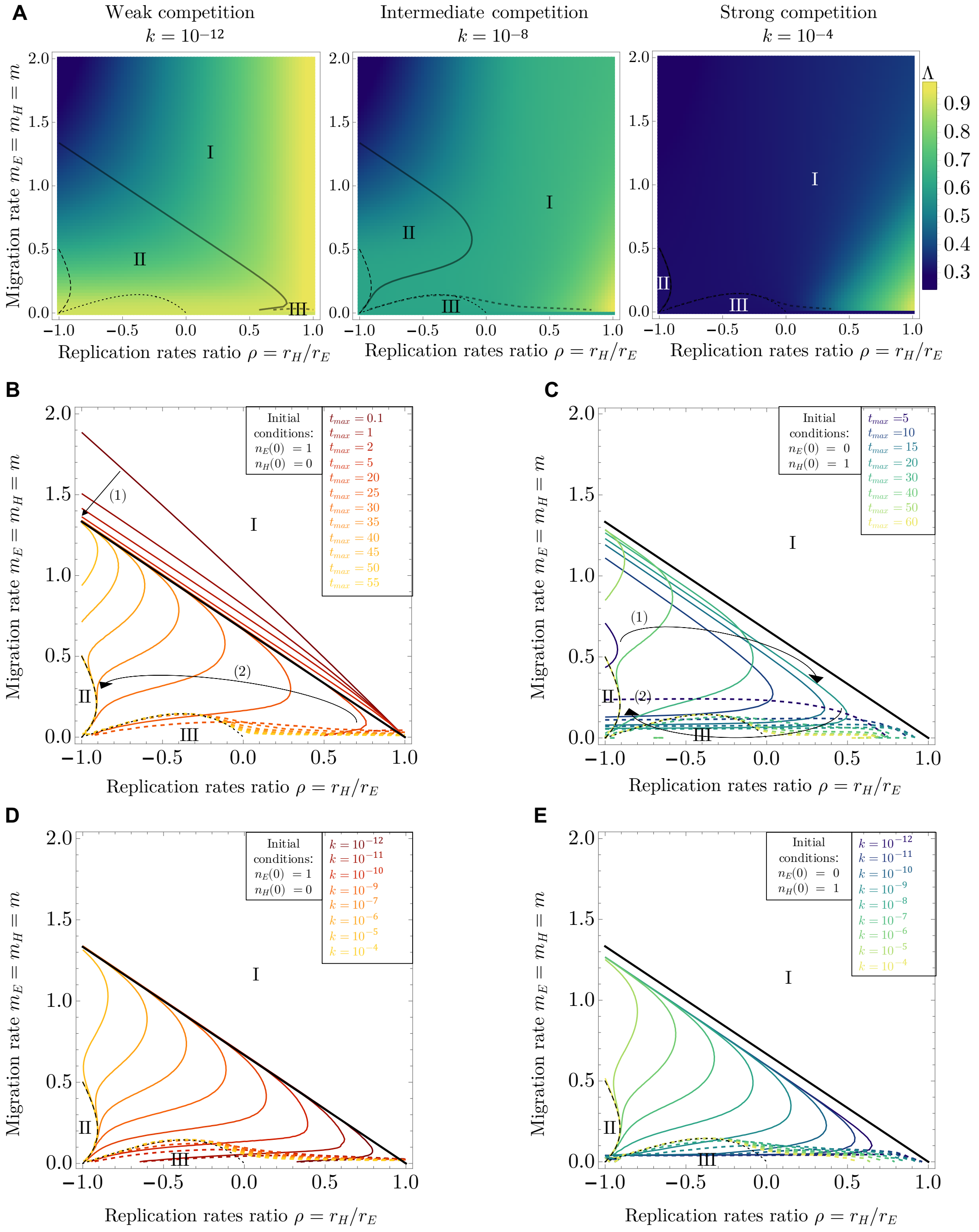


Figure A.2: **Optimal strategies in the model with limited growth in the environment only**. All the parameter values are identical to the ones in figure 4. **(A)** Change in the fitness landscape with $k$, the intensity of competition within the environment. The thick black lines show the contour line of equal sensitivities (solid: delimiting strategies I and II, dashed, strategies I and III). The thin black lines show the contours of equal sensitivities of the equilibrium population sizes (dashed: delimiting I and II, dotted: delimiting I and III).**(B-E)** Change in the contour lines delimiting the regions of optimality of the strategies with **(B-C)** $t_{max}$ (the time chosen to measure the final population size) and **(D-E)** $k$ (the intensity of within-host competition), in the case of initial conditions where all the microbes are in the environment (B and D, $n_E(0) = 1, n_H(0) = 0$) or in the case where all the microbes are initially in the host (C and E, $n_E(0) = 0, n_H(0) = 1$). The solid lines show the limit between the regions of optimality of strategies I and II (strategy I: increasing the replication rates ratio; strategy II: decreasing migration), and the dashed lines between the strategies I and III (strategy III: increasing migration). The thin black lines show the contours of equal sensitivities of the equilibrium population sizes (dashed: delimiting I and II, dotted: delimiting I and III). In the case of competition limited to the environment, whatever be the initial conditions, at sufficiently large times or intensity of competition, the strategy II (decreasing migration) tends to see its area of optimality drastically reduced.
